# The phosphoproteome of rice leaves responds to water and nitrogen supply

**DOI:** 10.1101/2021.03.15.435047

**Authors:** Sara Hamzelou, Vanessa J. Melino, Darren C. Plett, Karthik Shantharam Kamath, Arkadiusz Nawrocki, Martin R. Larsen, Brian J. Atwell, Paul A. Haynes

## Abstract

The scarcity of freshwater is an increasing concern in flood-irrigated rice, whilst excessive use of nitrogen fertilizers is both costly and contributes to environmental pollution. To co-ordinate growth adaptation under prolonged exposure to limited water or excess nitrogen supply, plants have processes for signalling and regulation of metabolic processes. There is limited information on the involvement of one of the most important post-translational modifications (PTMs), protein phosphorylation, on plant adaptation to long-term changes in resource supply. *Oryza sativa* cv. Nipponbare was grown under two regimes of nitrogen from the time of germination to final harvest. Twenty-five days after germination, water was withheld from half the pots in each nitrogen treatment and low water supply continued for an additional 26 days, while the remaining pots were well watered. Leaves from all four groups of plants were harvested after 51 days in order to test whether phosphorylation of leaf proteins responded to prior abiotic events. The dominant impact of these resources is exerted in leaves, where PTMs have been predicted to occur. Proteins were extracted and phosphopeptides were analysed by nanoLC-MS/MS analysis, coupled with label-free quantitation. Water and nitrogen regimes triggered extensive changes in phosphorylation of proteins involved in membrane transport, such as the aquaporin OsPIP2-6, a water channel protein. Our study reveals phosphorylation of several peptides belonging to proteins involved in RNA-processing and carbohydrate metabolism, suggesting that phosphorylation events regulate the signalling cascades that are required to optimize plant response to resource supply.

## 1. Introduction

Rice is a major crop cultivated globally, especially in developing countries in tropical and sub-tropical Asia. In spite of its importance as a prominent component of food security, productivity is vulnerable to erratic water supply in many systems of cultivation [1] and soil nitrogen status [2]. Stabilising production systems by optimizing plant response to variable resource supply will lead to more efficient use of resources, especially in low-input rainfed production systems. Water and nitrogen are critical resources for economic yields in all crops. Nitrogen is a core driver of plant productivity because it is a major component of amino acids, nucleotides, and some hormones and defence compounds [3]. However, acquisition of nitrogen is not independent of water supply, with rice having particularly high transpiration rates and, in cultivated systems, high nitrogen demand. In addition, under drought stress, rice plants treated with ammonium were shown to have higher water absorption than nitrate treated plants [4], and this response was correlated with the higher expression of aquaporin genes [5,6]. It has also been long known that transpiration directly affects nitrogen acquisition, with drought reducing the mass flow of soil nitrogen and delivery to leaves through the xylem, thereby compromising nitrogen assimilation and, ultimately, photosynthesis [7,8]. Knowing that these interactions exist, it is unsurprising that plants have evolved mechanisms to optimize the efficient use of resources in leaves, leading to the evolution of internal signalling pathways for both water [9] and nitrogen [10] economies, as well as the likelihood of crosstalk between these pathways [11]. The scale and sensitivity of gene expression responses to plant water and nitrogen status are indicative of the complex gene regulatory networks required to optimize resource use in plants [11].

Until quite recently, cell responses to environmental stresses have primarily been viewed as via regulation of metabolic cascades through remodelling the transcriptome and proteome. However, as well as the effects of stress on the abundance of proteins, post-translational modifications (PTMs) including phosphorylation are now widely acknowledged as important regulators of the catalytic activity, stability, and localization of proteins [12]. Specifically, plants respond to environmental cues through the regulation of signalling mechanisms mediated partially by protein PTMs [13]. Phosphorylation is one of the most important PTMs and is essential to a deeper understanding of drought stress signalling in plants [12]. Phosphoproteomic studies to date have focussed mostly on changes in response to short-term stressors such as dehydration, although long-term exposure to water stress has been investigated recently [14,15]. For example, comparative phosphoproteomic studies on short-term and long-term exposure of plants to osmotic stress showed some common phosphorylation events that are conserved over varying time scales [14,15]. Therefore, it was of interest to investigate alterations in the phosphoproteome in response to the supply of water and nitrogen. In that water and nitrogen deficits are strongly expressed in leaves, these experiments involved sampling vegetative tissue after plants had reached a steady-state over many weeks of treatment.

The low abundance of PTMs calls for efficient, sensitive methods to enrich them, partly accounting for the dearth of detail on PTMs in plants under stress [16]. Along with improvements in mass spectrometry instrumentation and developments in PTM enrichment methods, more research is required in the area of protein PTMs. Using titanium dioxide (TiO_2_) chromatography enrichment and subsequent nanoLC-MS/MS analysis, coupled with label-free quantification, we performed a detailed quantitative phosphoproteomic analysis of *O. sativa* (cv. Nipponbare) leaves grown in 40% and 100% field capacity (FC) at optimum (250 mg N / kg) and elevated (750 mg N / kg) nitrogen levels. These treatments were designed to reflect a range of resource regimes as commonly seen in the field, without inducing severe drought or nitrogen deficiencies and tissue senescence. This revealed novel insights into the impact of water and nitrogen regimes (and their interaction) on the phosphoproteome of *japonica* rice in late vegetative development.

## 2. Materials and methods

### Plant growth and sampling

Rice (*Oryza sativa* cv. Nipponbare) plants (*n* = 4) were grown in 40-cm cylindrical pots (16 cm diameter) containing 8.6 kg of soil consisting of 50% (v/v) University of California mix [50% (v/v) Waikerie river sand: 50% (v/v) peatmoss], 35% (v/v) peat mix and 15% (v/v) clay loam. Soil samples were dried at 80°C for 5 days with periodic weighing until there was no more loss in weight and gravimetric water content was calculated in g water per g soil. Ammonium nitrate was added in two doses to obtain a final concentration of 250 (N250) and 750 (N750) mg N per kg of soil in the respective treatments. One plant was grown per pot in a randomized complete block design with four replicates. The experiment was conducted in an automated gravimetric watering system (DroughtSpotter, Phenospex, Heerlen, The Netherlands) in a controlled environment room at the Plant Accelerator, Australian Plant Phenomics Facility, at the University of Adelaide, South Australia. Plants were grown in 12 h/12 h (light/dark) under a light intensity of 700 μmol m^−2^ s^−1^ at 30/22°C (day/night). The amount of water applied and the weight of pots was recorded automatically either hourly or half-hourly during the experiment using the DroughtSpotter [17]. From 25 - 51 days after planting (DAP), well-watered (WW) plants were watered to saturation, while water deficit (WD) plants were watered to 40% field capacity (FC). Flag leaves were harvested from all plants at 51 DAP between 12:00 and 13:00, weighed, and immediately frozen in liquid nitrogen, followed by lyophilization and grinding into a fine powder for protein extraction. Shoot and root biomass was measured after oven-drying at 70 °C to a constant weight.

### Gas exchange measurements

Transpiration rate (*T*_r_) and assimilation rate (*A*_r_), were measured at 51 DAP. Gas exchange was measured between 10:00 to 12:00 using a portable LI-COR photosynthesis system (LI-6800; LI-COR, Inc., Lincoln, NE, USA) with a pulse-amplitude modulated (PAM) leaf chamber head. Data were collected from the youngest fully expanded leaves with the following parameters: light intensity at 1800 µmol m^-2^ s^-1^, CO_2_ concentration of 400 µmol mol^-1^, leaf chamber temperature of ∼ 28°C, and relative humidity of ∼50%.

### Protein extraction, protein assay, and in-solution digestion

Protein extraction and in-solution digestion were performed as described previously [18]. In brief, 40 mg lyophilized leaf powder from all biological replicates was suspended in 1.5 ml of 10% (v/v) trichloroacetic acid in acetone, 2% (v/v) β-mercaptoethanol, followed by 30 min vortex at 4 °C and incubation at -20 °C for 45 min. As the leaf samples were simultaneously snap frozen using liquid nitrogen, which arrests the cellular activity, phosphatase and protease inhibitors were not used in the lysis buffer [19]. The pellet was collected after 16,000 × g centrifugation for 30 min and washed three times with ice-cold acetone. The resulting protein pellet was lyophilized in a vacuum centrifuge and resuspended in 3% (w/v) SDS in 50 mM Tris-HCl (pH 8.8), and then precipitated using methanol-chloroform. The resulting protein pellet was suspended in 8 M urea in 100 mM Tris-HCl, pH 8.8. Protein concentration was measured using bicinchoninic acid (BCA) assay kit (Thermo Scientific, San Jose, CA, USA). Protein samples were diluted five-fold using 100 mM Tris-HCl, (pH 8.8), reduced with 10 mM dithiothreitol at room temperature for 1 h, and alkylated with 20 mM iodoacetamide in the dark for 45 min at room temperature. Samples were digested at 37°C overnight using trypsin at an enzyme to protein ratio of 1:50. To stop the digestion, the pH was reduced to 3 or less with the addition of trifluoroacetic acid (TFA) to 1 % of the total volume.

### TiO_2_ phosphopeptide purification

Phosphopeptides were enriched using TiO_2_ chromatography according to Thingholm et al, with minor modifications [20]. Briefly, digested peptides were resuspended in loading buffer containing 80% acetonitrile (ACN), 5% TFA, and 1 M glycolic acid, followed by incubation with the corresponding amount of TiO_2_ (0.6 mg TiO_2_ beads per 100 μg peptides) for 15 min at room temperature with continuous mixing. The peptides and TiO_2_ beads mixture were centrifuged for 15 s. In order to increase the enrichment, the supernatant containing previously unbound peptide was incubated with 0.3 mg TiO_2_ beads per 100 μg peptides for 10 min with continuous mixing. The TiO_2_ beads were pooled and transferred to fresh low-binding microcentrifuge tubes. The beads were washed two times, first with 80% ACN, 1% TFA, and then with 10% ACN, 0.2% TFA. After drying the beads using a vacuum centrifuge for 10 min, the phosphopeptides were eluted from the beads by adding 1.5% ammonium hydroxide (pH 11.3) for 15 min. The samples were centrifuged at 13,000 rpm for 1 min at room temperature, followed by passing the eluate through a C8 stage tip eluted with 30% ACN. The peptides were lyophilized, and redissolved in 20 mM triethylammonium bicarbonate (TEAB) buffer (pH 7.5). For deglycosylation, Peptide /N-Glycosidase F (PNGaseF) was added at an enzyme: protein ratio of 1:150 and incubated at 37°C overnight. pH was reduced to less than 3 by adding TFA and, samples were desalted using Poros Oligo R3 stage tip [21]. Phosphopeptides were washed with 0.1% (v/v) formic acid and eluted by adding 70% (v/v) ACN, 0.1% (v/v) formic acid. The eluent was dried using a vacuum centrifuge and resolubilized in 0.1% (v/v) formic acid.

### Nanoflow liquid chromatography-tandem mass spectrometry

Phosphopeptide samples were analysed by nanoflow liquid chromatography-tandem mass spectrometry (nLC–MS/MS) using a Q Exactive Orbitrap mass spectrometer coupled to an EASY nLC1000 nanoflow HPLC system (Thermo Scientific, San Jose, CA, USA). Phosphopeptides were loaded on reversed phase columns of 75 µm internal diameter that were packed in-house to 10 cm with Halo C18 packing material (2.7 µm beads, 160 Å pore size, Advanced Materials Technology) [22]. The applied gradient for eluting the phosphopeptides consisted of mobile phase A (0.1% formic acid) and mobile phase B (99.9% (v/v) ACN, 0.1% (v/v) formic acid). A 100-min linear solvent gradient was used, starting with 2% mobile phase B for 1 min, 2 - 30% for 89 min, 30 - 85% for 5 min; and 85% B for another 5 min. Full MS scan range of 350 to 1850 *m*/*z* was acquired in the Orbitrap at resolution of 70,000 at *m*/*z* and automatic gain control (AGC) target value of 1 × 10^5^. Data-dependent MS/MS analysis of the ten most intense ions was conducted at 27% normalized collision energy and dynamic exclusion of ions for 20 s.

### Protein identification

Raw MS data files were processed with MaxQuant 1.6.12.0 loaded with Andromeda search engine [23] and searched against the rice (*Oryza sativa*) proteome fasta database downloaded from UniProt in April 2020 (48,904 protein sequences). Protein N-terminal acetylation, methionine oxidation, and phosphorylation on serine, threonine, and tyrosine (STY) were selected as variable modifications and cysteine carbamidomethylation was selected as a fixed modification. Trypsin was specified as the digesting enzyme, and up to two missed cleavages were allowed. The false discovery rate was set at 1% for proteins and peptides and matching between runs was performed within a time window of 1 min. The mass tolerance for full scans was set to 20 ppm for precursor and 0.5 Da for fragment ions. Label-free quantitation was performed using normalized label-free quantification (LFQ intensity) using the MaxLFQ algorithms as described [24].

### MS data processing and analysis

Intensities from “phospho(STY)sites.txt” file were used for further analysis. The intensity value of two technical replicates for each of three biological replicates was averaged when a phosphosite was quantified in both technical replicates. Data processing and statistical analysis of MaxQuant was performed in Perseus 1.6.0.2 [25]. Intensities were log_2_ transformed and high-confidence identification filtering was applied by maintaining phosphorylation events with a localization probability above 0.75 (referring to 75% confidence in localization of phosphorylation). The dataset was filtered based on phosphosites that were quantified in at least two out of three biological replicates of each group of nitrogen and water treatments. Missing values of quantification were imputed using normal distribution. A pairwise comparison of phosphosite quantification values between well-watered (WW) and water deficit (WD) plants grown in either N250 or N750 was performed using two-tailed *t*-tests. Differentially expressed phosphosites were identified based on *p*-value < 0.05.

### Physiological data statistical analysis

Data from physiological analysis of four biological replicates including shoot dry weight, daily water consumption, root to shoot ratio (R:S), and gas exchange measurements were processed using R (version 3.6.1) [26], with a two-way ANOVA. Differences between means were tested with a Tukey HSD *post hoc* test and *p*-value < 0.05 was considered significant.

### Functional protein annotation and motif analysis

Gene ontology (GO) annotations were obtained from the UniProt database and matched to the list of significantly changed phosphoproteins. Phosphoproteins were classified into functional categories of interest using PloGO [27]. The information related to sub-cellular localization was gained from an online prediction tool, Plant-mSubP [28]. Sequence motif analysis was performed using IceLogo [29] using the sequence windows for individual phosphosites acquired from MaxQuant, and the rice proteome as the background dataset. Results were presented using percent difference as a scoring system and a *p*-value cut-off of 0.05.

## 3. Results

### 3.1 Whole-plant effects of water and nitrogen regimes

The combination of adequate watering and an optimal nitrogen regime for 51-d-old plants (WW/N250) resulted in heavier leaf canopies and higher root-to-shoot ratios than observed in the other three treatments (Figure 1a). Shoot growth was slowed by withholding water (WD) but unaffected by nitrogen supply, with leaves in neither treatment exhibiting any symptoms of chlorosis or tissue death. This reflected the choice of nitrogen treatments, which were designed to fall within the optimal range for healthy growth. Reduced nitrogen supply increased the root-to-shoot ratio (R:S) in WW plants, as is typical for nitrogen-deficient plants [30], while N750 plants invested less than 20% of their biomass in roots (Figure 1a).

**Figure 1.**
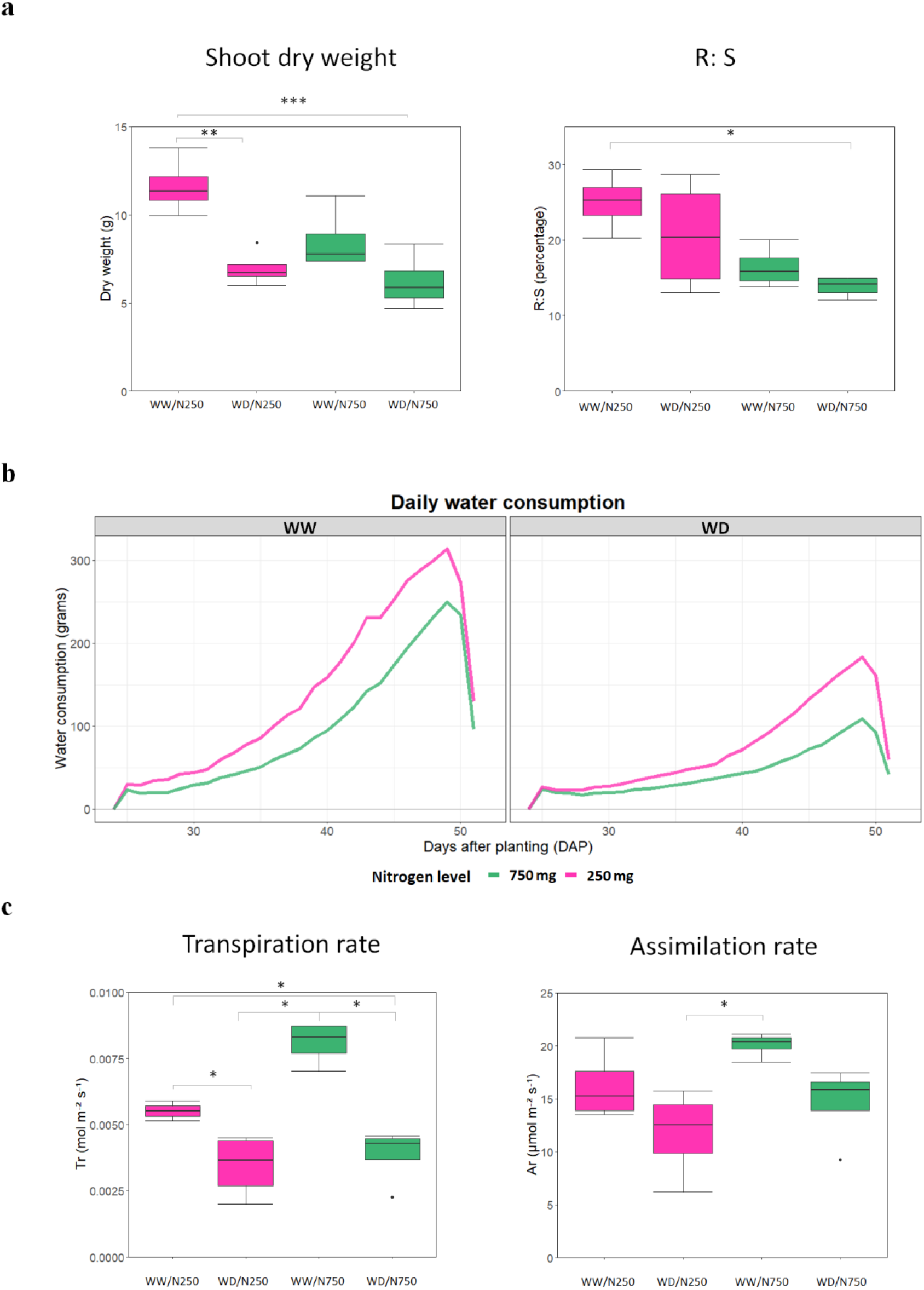
Growth, water consumption and leaf gas exchange in various water and nitrogen treatments. (**a)** shoot dry weight and root-to-shoot ratios. Asterisks show values that are statistically significantly different, according to ANOVA plus Tukey HSD *post hoc* test and *t*-student test. (**b)** water consumption in each of four replicates in WW, WD in N750 and N250 nitrogen from 25 to 51 DAP. (**c)** leaf gas exchange including transpiration rate (left), and assimilation rate (right). Asterisks indicate statistically differences between different treatments acquired from ANOVA plus Tukey HSD *post hoc* test.

Daily water consumption of whole plants rose steeply in the latter phase of the experiment and was consistently about 50% lower in the WD treatment (Fig. 1b), reflecting the impact of limited water supply. Curiously, water consumption was significantly higher in N250 plants compared with N750 plants, regardless of watering regime, as a result of the more rapid canopy expansion in the earliest stages of development in N250 plants (Fig. 1b). This is illustrated by transpiration rates *per unit of leaf area*, as measured by gas exchange techniques, where rates in N750 plants were equivalent or higher than those of N250 (Fig. 1c). Notably, transpiration rates (*T*_r_) were lower in WD plants compared with WW plants, whether they were grown at N250 or N750 (Figure 1c). Assimilation rates (*A*_r_) generally followed the same patterns as *T*_r_ but only the combination of higher levels of water and nitrogen supply resulted in significantly higher *A*_r_ than in the other three treatments (Figure 1c).

### 3.2 Phosphopeptides identified in leaves of plants grown in two nitrogen treatments under water deficit

In order to investigate the effects of soil nitrogen concentration during limited water deficit on global protein phosphorylation status, the changes in protein phosphorylation in leaf samples were analysed by comparing WD with WW plants grown in N250 and N750. Phosphoproteome analysis was performed using selective enrichment of phospho-peptides using TiO_2_ chemistry followed by nanoLC-MS/MS analysis in two technical replicates and three biological replicates. A total of 2232 phosphosites on 401 phosphoproteins were identified (Supplementary Table 1), of which 1200 phosphosites were localized with a probability of above 75%. A total of 309 phosphosites related to 250 phosphoproteins were identified in at least two replicates in one or more of the four treatments.

The majority of identified phosphosites were singly phosphorylated peptides (93%) and the remaining 7% were doubly phosphorylated (Figure 2a). Of the identified phosphosites, 80% were phosphorylated on serine, 19% on threonine, and only 1% on tyrosine (Figure 2b). The effects of water deficit are reported at the same level of nitrogen (WD/N750 vs. WW/N750; WD/N250 vs. WW/N250) as well as effects of the interaction between water and nitrogen supply (WD/N250 vs. WW/N750, and WD/N750 vs. WW/N250). Of all the phosphosites that were statistically significantly altered (*p*-value< 0.05) in relative abundance, phosphorylation of Thr881 on plasma membrane H^+^-ATPase (Q7XPY2) and Ser1077 were found to be decreased in drought stress in all four comparisons (Figure 3d).

**Figure 2.**
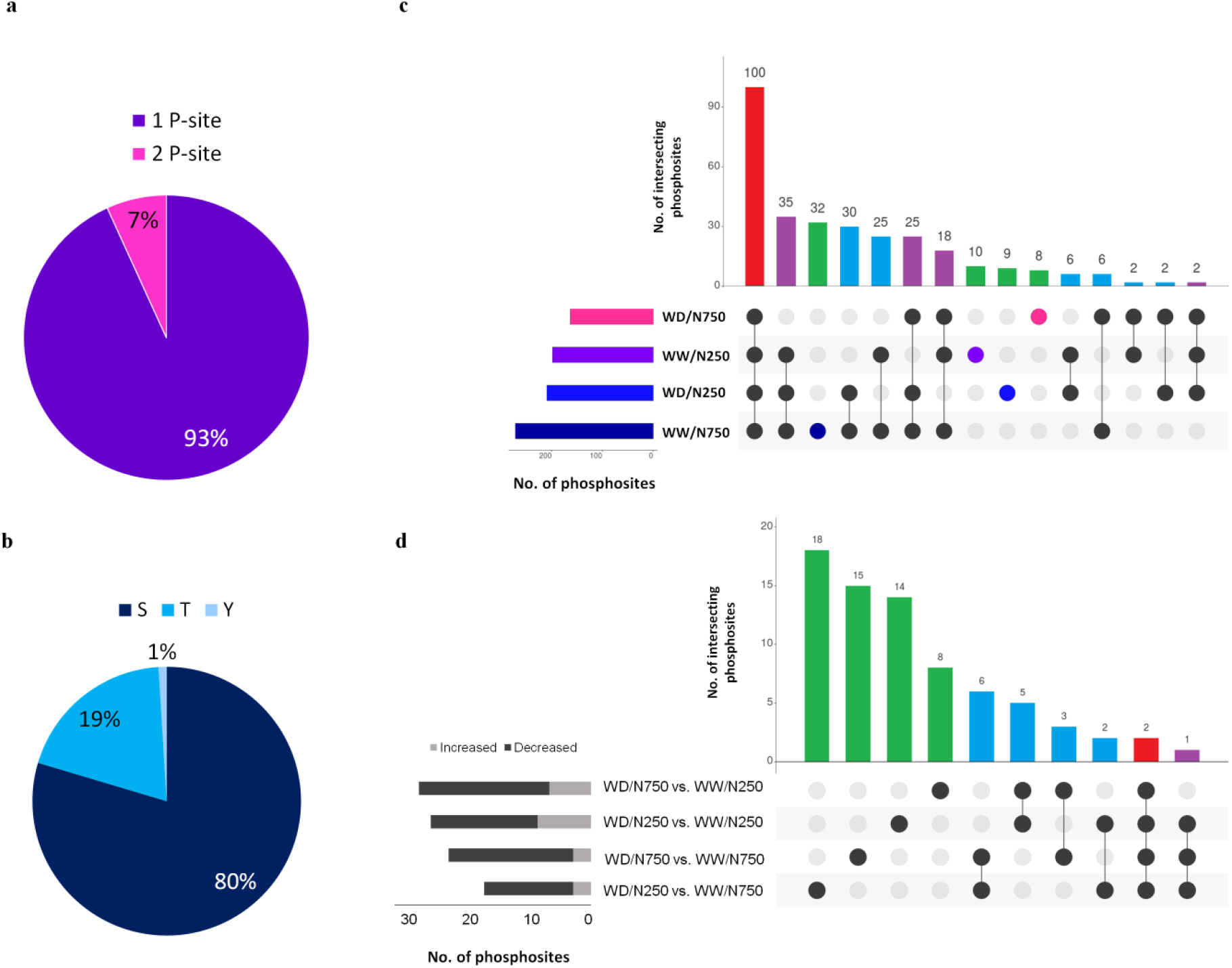
Characterisation and analysis of identified phosphosites. (**a**) Distribution of phosphorylation sites by number of phosphorylated sites. (**b**) Phosphorylated amino acid residue composition (**c**) UpSet visualisation [31] of the number of identified phosphosites (**d**), and significantly changed phosphosites in phosphorylation along with the number of intersecting phosphosites. Filled circles and vertical lines represent the corresponding phosphosites being compared. left bar graph for the UpSet plot d shows the number of significantly increased (light gray) and decreased (dark gray) phosphorylation changes in each of four comparisons upon water deficit and nitrogen supply.

**Figure 3.**
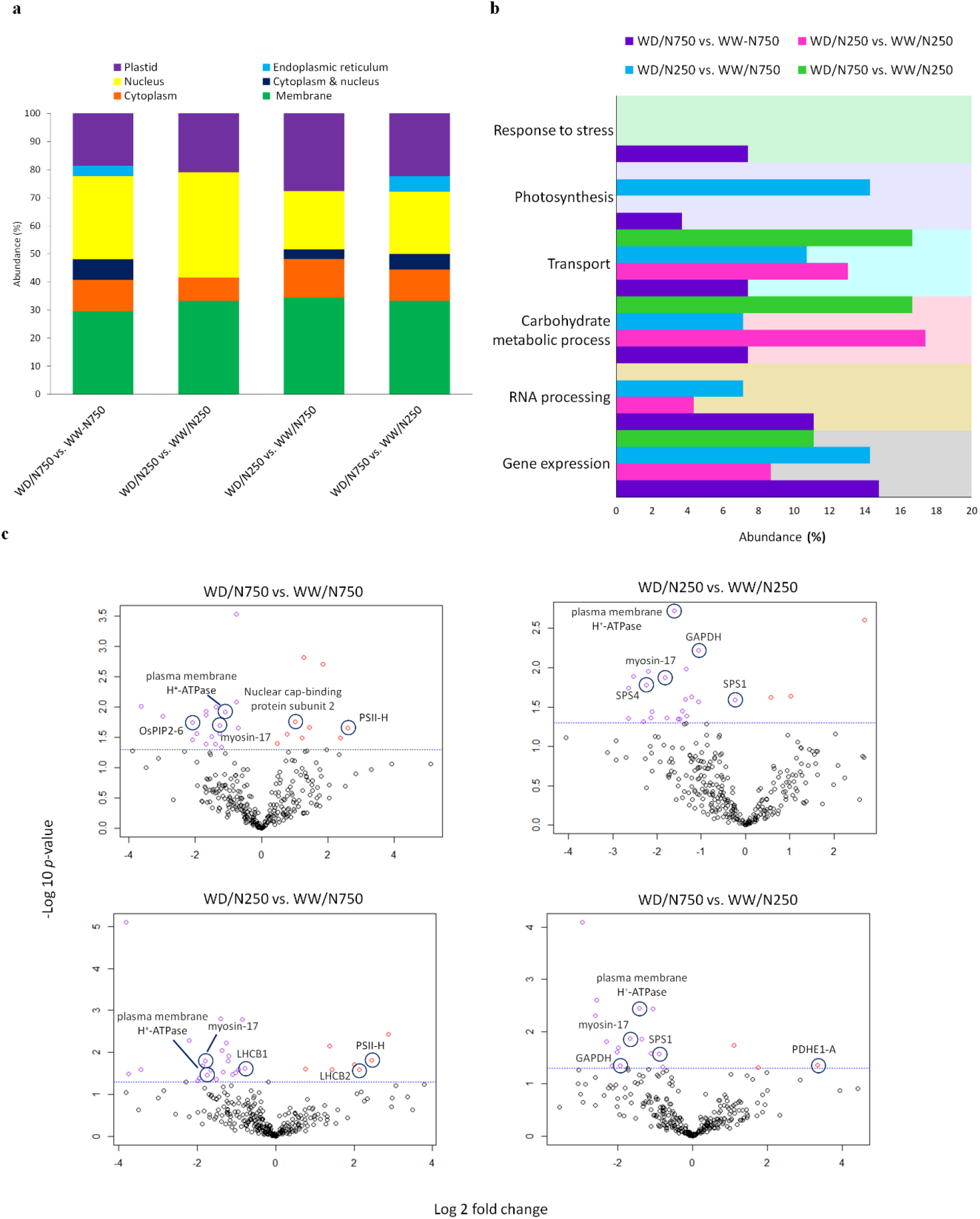
Functional classification and subcellular localization of differentially phosphorylated phosphoproteins under water stress and nitrogen treatment. (**a)** Subcellular localization of significantly changed phosphoproteins in different water and nitrogen treatments. (**b)** Functional classification of differentially phosphorylated phosphoproteins under drought and different nitrogen treatments. (**c)** Volcano plots of all significantly changed phosphoproteins in each of four comparisons. Each point illustrates a phosphoprotein with an average log 2-fold change along the x-axis and –log10 *p*-value along the y-axis. Red, purple, and black points indicate the increased, decreased, and unchanged phosphorylation, respectively. The dashed blue line shows the *p*-value of 0.05 cut-off.

### 3.3 Functional Analysis and Subcellular Localization of phosphosites upon water and nitrogen regimes

Subcellular localization and biological processes enriched in significantly altered phosphoproteins of different comparisons were analysed (Figure 3). The identified phosphoproteins were predominantly localized in either cell membrane or nucleus (Figure 3a). Only two proteins were significantly changed in phosphorylation in all four comparisons; myosin-17 and plasma membrane H^+^-ATPase. The most abundant functional categories of the differentially phosphorylated proteins were gene expression and carbohydrate metabolic processes. Of those within the gene expression functional group, RNA processing and mRNA metabolic processing (i.e., mRNA splicing) were the major categories (Figure 3b). Further information on the main functional categories of phosphoproteins for each of four comparisons is presented below.

#### 3.3.1 Plant response to water deficit in N750

Out of 282 identified phosphosites in this comparison group, 27 phosphosites, belonging to the same number of phosphoproteins, were significantly changed in phosphorylation in response to water stress. Eight of these (30%) were membrane proteins, including probable aquaporin PIP2-6 (Q7XLR1). The phosphosite MS(p)KEVSEEPEHVRPKDY from that protein was phosphorylated at ser2 in all replicates of well-watered plants grown in N750, but only detected at low level in one replicate of the water-deficit plants. A significant change in phosphorylation of this peptide in PIP2-6 was observed in both this comparison and WD/N250 vs. WW/N750. Gene expression-related phosphoproteins, especially those involved in RNA processing, were found to be one of the most abundant functional categories in this group. These proteins included La protein 1, Hepatocellular carcinoma-associated antigen 59 family protein, and Nuclear cap-binding protein subunit 2. Interestingly, this comparison was the only one in which stress-responsive phosphoproteins were found to be significantly changed in phosphorylation (Figure 3b). These included Nodulin-related protein 1 (NRP1) and EMSY-LIKE 3 protein that displayed changes in phosphorylation of Ser9, and Ser477, respectively, following the imposition of drought stress in N750.

#### 3.3.2 Plant response to water deficit in N250

A total of 290 phosphosites were detected to be altered upon treatment of rice plants with drought stress and low nitrogen exposure, of which 24 phosphosites corresponding to 23 phosphoproteins were significantly changed in abundance of phosphosites. This included nine nucleus localised phosphoproteins and eight membrane localised proteins that were significantly changed in phosphorylation. Two of the significantly changed nucleus localised phosphoproteins, Q0E0T0 and Q69XK9, were annotated as uncharacterised proteins. Four of the significantly changed phosphoproteins were involved in glycolysis and sucrose biosynthetic processes. Those involved in sucrose biosynthesis include probable sucrose-phosphate synthase 1 (SPS1) and probable sucrose-phosphate synthase 4 (SPS4). Phosphorylation of Ser179 and Ser170 residues were decreased significantly in SPS1 and SPS4, respectively, in response to water deficit for the plants treated with N250. Ser289 of glyceraldehyde-3-phosphate dehydrogenase (GAPDH) and Ser205 of glyceraldehyde-3-phosphate dehydrogenase isoform 2 were also decreased in phosphorylation.

#### 3.3.3 Plant response to water deficit from N750 to N250 (WD/N250 vs. WW/N750)

From 290 identified phosphosites in this comparison group, 29 phosphosites corresponding to 28 phosphoproteins were significantly changed in phosphorylation. Although a significant number of phosphoproteins were localized in the chloroplast in all four comparisons, phosphoproteins in the photosynthesis functional category were enriched mostly in this comparison group (Figure 3a). However, Photosystem II reaction center protein H (PSII-H) was significantly changed in phosphorylation not only in this comparison group but also in those grown in N750.

Chlorophyll a-b binding protein 1 (LHCB1), Chlorophyll a-b binding protein 2 (LHCB2), and PGR5-like protein 1A were identified to have significantly altered phosphorylation only in this comparison group. Though LHCB1 and LHCB2 are members of LHCB family, the alteration in phosphorylation patterns in these proteins differed. Phosphorylation of Thr35 was significantly decreased in LHCB1 while phosphorylation of Ser32 was significantly increased in LHCB2 in response to water stress (Figure 3c).

#### 3.3.4 Plant response to water deficit from N250 to N750 (WD/N750 vs. WW/N250)

Out of 197 identified phosphosites in this comparison group, 18 phosphosites from 18 phosphoproteins were significantly changed in phosphorylation. These phosphoproteins were categorized mainly in three GO categories: gene expression, carbohydrate metabolic processes, and transport (Figure 3b). Interestingly, of those phosphoproteins involved in carbohydrate metabolic processes, pyruvate dehydrogenase E1 component subunit alpha-1 was found to be increased in phosphorylation in response to water deficit only in this group. SPS1 and GAPDH were also decreased in phosphorylation, in agreement with the findings in plant response to water stress in N250 (Figure 3c).

Along with myosin-17 and plasma membrane H^+^-ATPase, that were identified in all four groups, putative mannitol transporter was significantly decreased in phosphorylation at ser250 in response to water deficit. Greatly increased phosphorylation of putative mannitol transporter in well-watered plants in N250 resulted in a significant change in phosphorylation of this protein when compared to water-deficit plants in N250, and also in N750.

### 3.4 Motif analysis of significantly changed phosphosites

Linear motif analysis was performed for significantly changed phosphosites in four comparisons. In general, there was a strong representation of proline and arginine in proximity to the phosphorylated serine and threonine residues. Arginine residue (R) at the -3 position (Rxx(S/T)) was enriched for all comparisons. However, proline-directed motif ((S/T)/P) was also enriched, but to a lesser extent, in all comparisons apart from WD/N750 vs. WW/N250. Along with the basic Rxx(S/T) motif, glutamic acid was significantly over-represented at position +2 ((S/T)xxE) following drought response in N750. The enriched sequence motifs in each of the treatments are shown in figure 4.

**Figure 4.**
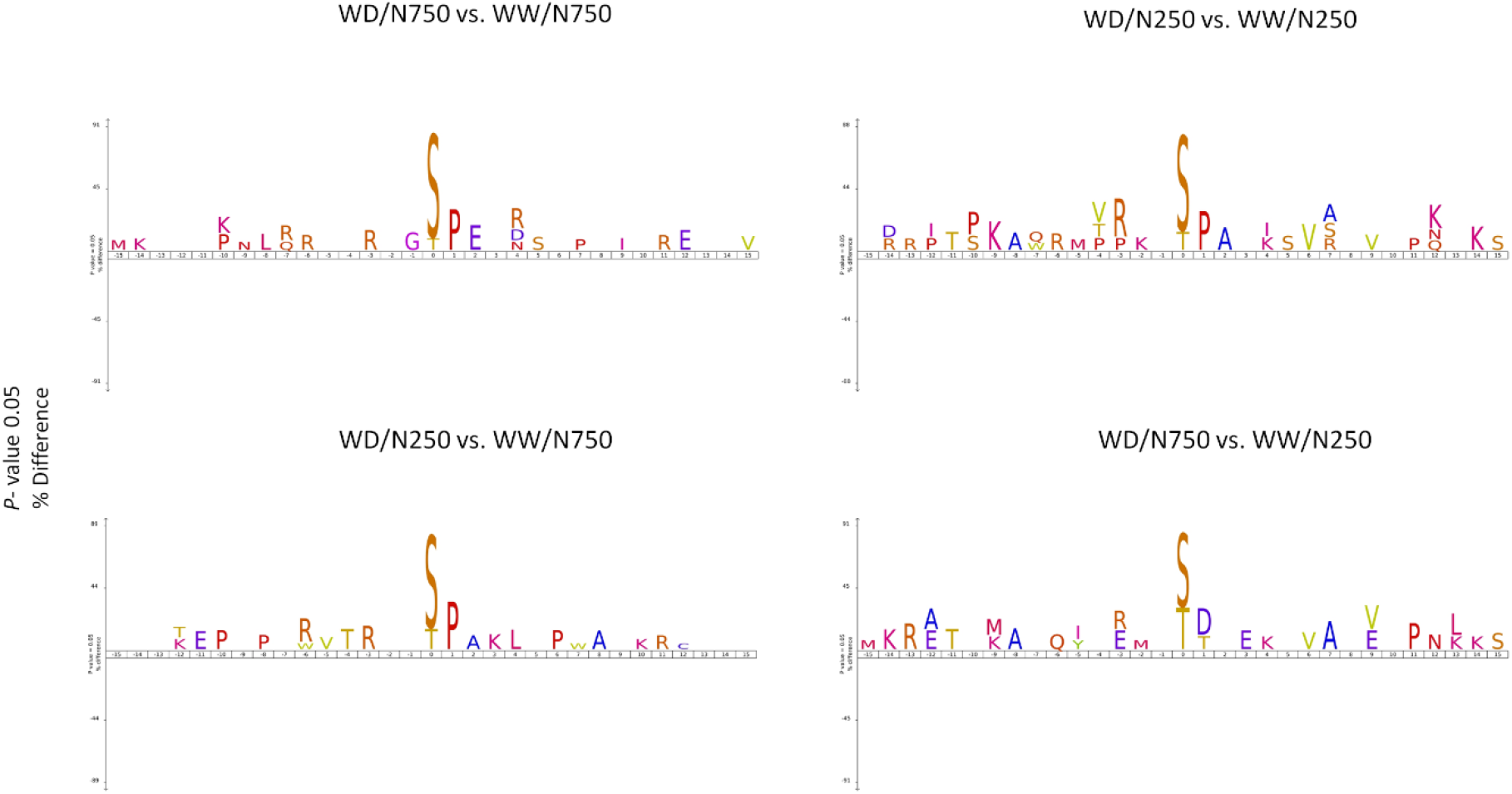
Motif analysis of changing phosphorylated phosphosites. The characters above the horizontal axis show the significantly enriched amino acids. (*p* < 0.05). The phosphorylated amino acid residues (S/T) are located at position 0.

## 4. Discussion

### 4.1 Physiological response of plants to water and external nitrogen

Sustained water deficits are a key inhibitor of rice growth [32] and inhibit assimilation rates, ultimately impairing yields [33]. Sustained nitrogen supply and water availability are essential resources for optimal plant growth and photosynthesis [3]. Previous studies on upland and lowland rice have indicated a positive correlation between optimum nitrogen application with aboveground biomass alterations [34,35]. In the current experiment, water supply was shown to limit plant performance independently of nitrogen supply, with WD plants unable to respond to the higher level of nitrogen (N750). Providing water was supplied at high levels (WW) by continuously monitored applications in the Plant Accelerator, the lower nitrogen treatment (N250) was sufficient to achieve healthy leaf canopies after only 51 d, even though assimilation rates were not significantly affected by nitrogen application (N250 *vs* N750) at the time of the final harvest. It has been shown that adequate nitrogen supply reduces water loss and enhances assimilation rate, indicating improved instantaneous transpiration efficiency [3]. High assimilation rates in WW plants at N750 compared with WD plants at N250 underline the significance of water and nitrogen as primary drivers of plant performance.

### 4.2 Effects of water and external nitrogen on phosphorylation alterations of transporter proteins

Based on the localisation of the majority of phosphoproteins in the cell and plastid membrane, phosphorylation appears to play a functional role in regulation of membrane proteins, especially transporters, in plants exposed to long-term water deficit and external nitrogen supply. Dephosphorylation of channel proteins was detected in the WD rice plants, which is mostly in correlation with inhibition of their activity for those well-characterized phosphoproteins such as plasma membrane H^+^-ATPase and aquaporin PIP2-6 [36–38]. The regulation of closure of aquaporin through phosphorylation has been previously linked to modulation of water loss in the leaf [39].

Those common phosphoproteins with differential phosphorylation levels in response to water stress in both nitrogen treatments such as plasma membrane H^+^-ATPase (Q7XPY2) and myosin-17 (A0A0P0W1J8), as well as a putative mannitol transporter, may reflect the dominant effect of water on the activity of such proteins. In contrast, the proteins found to be differentially phosphorylated in response to water supply at N750, such as aquaporin PIP2-6, may explain the impact of nitrogen on phosphorylation regulation of such channel proteins and the further regulation of water fluxes in plant cells.

Plasma membrane H^+^-ATPase proteins generate a proton gradient using the chemical energy of ATP, involving many physiological processes e.g. nutrient uptake, cell expansion, and stomatal opening [36]. PTMs modulate the activity of plasma membrane H^+^-ATPase, and phosphorylation at Thr-881 has been shown to activate this protein [36]. Based on phosphorylation alteration at Thr-881 of plasma membrane H^+^-ATPase, the activity of the protein declined in response to water stress in rice plants.

A significant difference was observed in phosphorylation of putative mannitol transporter in WW/N250 in comparison with WD plants in both N250 and N750. This membrane transport protein may play a role in photosynthetic carbon partitioning to mannitol which is an osmoregulator in plant cells [40]. Phosphorylation of myosin-17 and putative mannitol transporter may need to be investigated in further studies to address whether phosphorylation induces or inhibits the activity of these proteins. Previous studies have shown the accumulation of mannitol transporters in response to salt and osmotic stress in olive [40] and myosin-17 upon the pathogenic fungal stress in Arabidopsis [41].

PIPs (Plasma membrane Intrinsic Proteins) are a group of aquaporin channel proteins that act as gateways regulating the passage of water [38,42]. PTMs affect the activity of aquaporins through water flow modulation across the plasma membrane [3]. The phosphorylation of PIP channels is considered as the key mechanism for cell water flow regulation in response to environmental clues [43]. The phosphorylation of two serine residues on AtPIP2.1 was recently demonstrated to regulate the activity of this channel to shift between passage of water and ions [44]. Higher water absorption of rice grown in ammonium is connected with up-regulation of aquaporin genes [4,6]. The activation of aquaporins is regulated by signal transduction and PTM changes with respect to nitrogen amounts in shoot and root [3]. Our results confirmed the phosphorylation of probable aquaporin PIP2-6 in well-watered rice plants grown in N750, but not N250. This finding may explain the effect of external nitrogen and PTM on PIP2-6 activity, regarding the evidence related to the role of aquaporins in response to changes in nitrogen supply [3].

### 4.3 How do nitrogen and water deficit change the phosphorylation of RNA processing proteins?

Another group of phosphoproteins involved in RNA processing and mRNA metabolic processing was altered in response to water deficit. This finding is consistent with the fact that proteins involved in mRNA metabolism have been previously shown to contribute to environmental stress signalling that can be regulated by phosphorylation changes [45]. Importantly, the phosphoproteins in this group mainly play a role in the regulation of signalling activity of the plant hormone Abscisic acid (ABA). Serine/arginine-rich splicing factor (SR45 isoform X1), which decreased in phosphorylation level in WD/N250 plants in comparison with WW plants either in N250 or N750, plays a role in the regulation of pre-mRNA splicing. Dephosphorylation of SR proteins is required for their activity, such as interaction with RNA and splicing factors [46]. Thus, our results are predictive of an increased activity of SR45 under conditions of water stress and optimum nitrogen supply (N250). In contrast, 20 kDa nuclear cap-binding protein (CBP20) was increased in phosphorylation in water-deficit plants in both N250 and N750. CBP20 is one of the core proteins of the cap-binding protein complex, which is a contributor to different cellular mechanisms including pre-mRNA alternative splicing and those involved in growth, development, and response to environmental factors such as drought stress [47,48]. The activity of CBP20 may be increased by phosphorylation [49], so our results predict that CBP20 activity was induced under water deficit in a nitrogen-independent manner.

### 4.4 Carbohydrate signalling in response to changing nitrogen and water deficit

Plants respond to drought stress through carbohydrate signalling pathways and sugar sensing mechanisms. Soluble sugars are able to behave as signals regulating plant growth and development [50]. In the comparison of well-watered and droughted plants grown in N250, two isomers from SPS proteins, and two from GAPDH, were found to be differentially phosphorylated. GAPDH not only acts as a crucial enzyme in glycolysis but also plays important roles in various biological processes including plant response to environmental stresses. It has been demonstrated from a loss of function study that GAPDH mediates water loss [51]. In a recent study, the nuclear moonlighting of GAPDH has been suggested to be involved in conferring better tolerance of Arabidopsis to heat stress [52]. Phosphorylation changes were proposed as the main initial factor for stress-dependent translocation of this enzyme to the nucleus [53]. Carbohydrate metabolism may regulate plant response to drought through dephosphorylation of proteins involved in carbohydrate metabolic pathways [54]. *In vivo* phosphorylation of GAPDH at Ser205 results in a significant decrease in the activity of this enzyme during the seed development of wheat [55]. In agreement with previous studies, we observed that phosphorylation of GAPDH2 (Q7FAH2) at Ser205 significantly declined in droughted plants in comparison with well-watered plants grown in a lower amount of external nitrogen.

SPS plays a key role in sucrose synthesis and contributes to regulation of carbon assimilation and partitioning [56]. SPS was reported to be inhibited by phosphorylation in spinach leaves under osmotic stress [57]. It was also demonstrated that the phosphorylation level of SPS decreased significantly in wheat flag leaves under high nitrogen fertilizer [58]. We can postulate a relationship between dephosphorylation of GAPDH, along with SPS isoforms 1F and 4F, and changes in their activity that may contribute to the plant response to drought stress, especially in plants grown in N250.

### 4.5 Thylakoid membrane proteins are differentially phosphorylated in response to water and nitrogen regimes

Changes in phosphorylation of thylakoid membrane proteins play an important role in plant response to environmental stresses [59,60]. The change in phosphorylation events in thylakoid membrane proteins was mainly observed in WD/N250 vs. WW/N750. Although there is high sequence similarity between LHCB1 and LHCB2, these two members of the LHCB family have contrasting patterns of phosphorylation in response to stress. This is in agreement with a previous report showing different phosphorylation levels in LHCB1 and LHCB2 in response to changing light [61]. Photosystem II reaction center protein H (psbH) was also significantly changed in phosphorylation in water-deficit plants, either in N250 or N750. The significant increase in psbH concurs with a recent study that has shown the contribution of PbsH phosphorylation to PSII repair after photo-inhibition [62]. Abiotic stress causes PSII to become more vulnerable to photoinhibition [63], so repair of PSII seems to be required for plant viability.

### 4.6 Significantly over-represented sequence motifs in response to water and nitrogen

Proline-directed phosphorylation is one of the most common phosphorylation motifs in plants, and such proteins play a role in hormone signalling pathways such as ABA metabolism [64]. (S/T)/P motifs can be recognized by mitogen-activated protein kinases (MAPKs) [65]. MAPKs are involved in signalling pathways in response to multiple environmental stressors and developmental mechanisms [66,67]. MAPK signalling cascade is composed of three main kinases, MAP kinase kinase kinase (MAPKKK), that phosphorylates MAP kinase kinase (MAPKK), which finally phosphorylates MAPK [66]. The RxxS motif is also a potential substrate of MAPKK and subclass III SNF1-related protein kinase 2 (SnRK2), which is a major regulator in water-deficit conditions [68,69]. SxxE motif enrichment was also seen in the phosphosites targeted by casein kinase II (CKII). CKII is also engaged in signalling pathways activated by stresses [70]. All of the kinase motifs seen in our results, including (S/T)/P, RxxS/T, and SxxE, are known to be involved in cellular signalling cascades in plants in response to environmental stressors [65].

## 5. Conclusions

In this study, physiological response and PTM changes at the phosphorylation level of rice were studied in response to water deficit in the presence of varying nitrogen supplementation. In summary, the physiology data showed that there are essentially no statistically significant differences in measured parameters when nitrogen is varied but water status is kept constant, while, in contrast, there are statistically significant differences in the measured parameters when nitrogen is kept constant but plants are subjected to different watering regimes. Hence, we focused on the investigation of phosphorylation changes occurring in response to prolonged water deficit. Differential phosphorylation of membrane proteins, especially transporters, occurred in either a nitrogen-dependent or independent manner. This suggests that the PTM changes of transporter proteins may regulate water maintenance in leaf cells. RNA processing proteins were targeted by phosphorylation, indicating the importance of phosphorylation in the regulation of these proteins in response to drought. Our findings warrant further investigation regarding whether phosphorylation or dephosphorylation, is essential for the activation of these transporters and RNA processing proteins.

The phosphoproteins identified in this study, along with the quantitative information provided concerning changing levels of phosphorylation between different conditions, offer a resource for researchers and new insights into regulatory networks in plants in response to changing water and nutrient regimes.

## Abbreviations

ABA: Abscisic acid
ACN: Acetonitrile
A_r_: Assimilation rate
DAP: Days after planting
FC: Field capacity
GO: Gene ontology
LFQ: Label-free quantitation
N: Nitrogen
nLC–MS/MS: Nanoflow liquid chromatography-tandem mass spectrometry
PIP: Plasma membrane Intrinsic Protein
PTM: Post-translational modification
R:S: root-to-shoot ratio
T_r_: Transpiration rate
TFA: Trifluoroacetic acid
WD: Water deficit
WW: Well-watered

## Acknowledgments

SH would like to acknowledge scholarship support from Australian Commonwealth Government - International Research Training Program scholarship (iRTP). This work was supported by Macquarie University, and aspects of this research were conducted at the Australian Proteome Analysis Facility.

## References

1. Tuong, P., Bouman, B., Mortimer, M. More Rice, Less Water—Integrated Approaches for Increasing Water Productivity in Irrigated Rice-Based Systems in Asia. Plant Prod. Sci. 2005, 8, 231–241.

2. Gholizadeh, A., Saberioon, M., Borůvka, L., Wayayok, A., Mohd Soom, M.A. Leaf chlorophyll and nitrogen dynamics and their relationship to lowland rice yield for site-specific paddy management. Inf. Process. Agric. 2017, 4, 259–268.

3. Plett, D.C., Ranathunge, K., Melino, V.J., Kuya, N., Uga, Y., Kronzucker, H.J. The intersection of nitrogen nutrition and water use in plants: new paths toward improved crop productivity. J. Exp. Bot. 2020.

4. Gao, Y., Li, Y., Yang, X., Li, H., Shen, Q., Guo, S. Ammonium nutrition increases water absorption in rice seedlings (Oryza sativa L.) under water stress. Plant Soil 2010, 331, 193– 201.

5. Ding, L., Lu, Z., Gao, L., Guo, S., Shen, Q. Is Nitrogen a Key Determinant of Water Transport and Photosynthesis in Higher Plants Upon Drought Stress? Front. Plant Sci. 2018, 9, 1143.

6. Ding, L., Gao, C., Li, Y., Li, Y., Zhu, Y., Xu, G., Shen, Q., Kaldenhoff, R., Kai, L., Guo, S. The enhanced drought tolerance of rice plants under ammonium is related to aquaporin (AQP). Plant Sci. 2015, 234, 14–21.

7. Aragon, E.L., De Datta, S.K. Drought response of rice at different nitrogen levels using line source sprinkler system. Irrig. Sci. 1982, 3, 63–73.

8. Menge, D.M., Onyango, J.C., Yamauchi, A., Kano-Nakata, M., Asanuma, S., Thi, T.T., Inukai, Y., Kikuta, M., Makihara, D. Effect of nitrogen application on the expression of drought-induced root plasticity of upland NERICA rice. Plant Prod. Sci. 2019, 22, 180–191.

9. Takahashi, F., Shinozaki, K. Long-distance signaling in plant stress response. Curr. Opin. Plant Biol. 2019, 47, 106–111.

10. Xuan, W., Beeckman, T., Xu, G. Plant nitrogen nutrition: sensing and signaling. Curr. Opin. Plant Biol. 2017, 39, 57–65.

11. Araus, V., Swift, J., Alvarez, J.M., Henry, A., Coruzzi, G.M. A balancing act: how plants integrate nitrogen and water signals. J. Exp. Bot. 2020, 71, 4442–4451.

12. Wu, X., Gong, F., Cao, D., Hu, X., Wang, W. Advances in crop proteomics: PTMs of proteins under abiotic stress. Proteomics 2016, 16, 847–865.

13. Hashiguchi, A., Komatsu, S. Posttranslational Modifications and Plant–Environment Interaction. Methods Enzymol. 2017, 586, 97–113.

14. Bhaskara, G.B., Nguyen, T.T., Yang, T.-H., Verslues, P.E. Comparative Analysis of Phosphoproteome Remodeling After Short Term Water Stress and ABA Treatments versus Longer Term Water Stress Acclimation. Front. Plant Sci. 2017, 8, 523.

15. Bhaskara, G.B., Wen, T.-N., Nguyen, T.T., Verslues, P.E. Protein Phosphatase 2Cs and Microtubule-Associated Stress Protein 1 Control Microtubule Stability, Plant Growth, and Drought Response. Plant Cell 2017, 29, 169 LP – 191.

16. Arsova, B., Watt, M., Usadel, B. Monitoring of Plant Protein Post-translational Modifications Using Targeted Proteomics. Front. Plant Sci. 2018, 9, 1168.

17. Cousins, O.H., Garnett, T.P., Rasmussen, A., Mooney, S.J., Smernik, R.J., Brien, C.J., Cavagnaro, T.R. Variable water cycles have a greater impact on wheat growth and soil nitrogen response than constant watering. Plant Sci. 2020, 290, 110146.

18. Hamzelou, S., Kamath, K.S., Masoomi-Aladizgeh, F., Johnsen, M.M., Atwell, B.J., Haynes, P.A. Wild and Cultivated Species of Rice Have Distinctive Proteomic Responses to Drought. Int. J. Mol. Sci. 2020, 21, 5980.

19. Humphrey, S.J., Karayel, O., James, D.E., Mann, M. High-throughput and high-sensitivity phosphoproteomics with the EasyPhos platform. Nat. Protoc. 2018, 13, 1897–1916.

20. Thingholm, T.E., Larsen, M.R. The Use of Titanium Dioxide for Selective Enrichment of Phosphorylated Peptides. Methods Mol. Biol. 2016, 1355, 135–146.

21. Haynes, P.A. Phosphoglycosylation: a new structural class of glycosylation? Glycobiology 1998, 8, 1–5.

22. Thompson, E.L., Taylor, D.A., Nair, S. V; Birch, G., Haynes, P.A., Raftos, D.A. A proteomic analysis of the effects of metal contamination on Sydney Rock Oyster (Saccostrea glomerata) haemolymph. Aquat. Toxicol. 2011, 103, 241–249.

23. Cox, J., Mann, M. MaxQuant enables high peptide identification rates, individualized p.p.b.-range mass accuracies and proteome-wide protein quantification. Nat. Biotechnol. 2008, 26, 1367–1372.

24. Cox, J., Hein, M.Y., Luber, C.A., Paron, I., Nagaraj, N., Mann, M. Accurate proteome-wide label-free quantification by delayed normalization and maximal peptide ratio extraction, termed MaxLFQ. Mol. Cell. Proteomics 2014, 13, 2513–2526.

25. Tyanova, S., Temu, T., Sinitcyn, P., Carlson, A., Hein, M.Y., Geiger, T., Mann, M., Cox, J. The Perseus computational platform for comprehensive analysis of (prote)omics data. Nat. Methods 2016, 13, 731.

26. R Core Team. R: A language and environment for statistical computing. R Foundation for Statistical Computing, Vienna, Austria. 2012, ISBN 3-900051-07-0, URL http://www.R-project.org/.

27. Pascovici, D., Keighley, T., Mirzaei, M., Haynes, P.A., Cooke, B. PloGO: Plotting gene ontology annotation and abundance in multi-condition proteomics experiments. Proteomics 2012, 12, 406–410.

28. Sahu, S.S., Loaiza, C.D., Kaundal, R. Plant-mSubP: a computational framework for the prediction of single- and multi-target protein subcellular localization using integrated machine-learning approaches. AoB Plants 2020, 12.

29. Colaert, N., Helsens, K., Martens, L., Vandekerckhove, J., Gevaert, K. Improved visualization of protein consensus sequences by iceLogo. Nat. Methods 2009, 6, 786–787.

30. Lynch, J.P. Root phenotypes for improved nutrient capture: an underexploited opportunity for global agriculture. New Phytol. 2019, 223, 548–564.

31. Lex, A., Gehlenborg, N., Strobelt, H., Vuillemot, R., Pfister, H. UpSet: Visualization of Intersecting Sets. IEEE Trans. Vis. Comput. Graph. 2014, 20, 1983–1992.

32. Zhang, J., Zhang, S., Cheng, M., Jiang, H., Zhang, X., Peng, C., Lu, X., Zhang, M., Jin, J. Effect of Drought on Agronomic Traits of Rice and Wheat: A Meta-Analysis. Int. J. Environ. Res. Public Health 2018, 15.

33. Zhao, W., Liu, L., Shen, Q., Yang, J., Han, X., Tian, F., Wu, J. Effects of Water Stress on Photosynthesis, Yield, and Water Use Efficiency in Winter Wheat. Water 2020, 12.

34. Suriyagoda, L., Sirisena, D., Kekulandara, D., Bandaranayake, P., Samarasinghe, G., Wissuwa, M. Biomass and nutrient accumulation rates of rice cultivars differing in their growth duration when grown in fertile and low-fertile soils. J. Plant Nutr. 2020, 43, 251–269.

35. Wang, Y., Zhu, B., Shi, Y., Hu, C. Effects of Nitrogen Fertilization on Upland Rice based on Pot Experiments. Commun. Soil Sci. Plant Anal. 2008, 39, 1733–1749.

36. Falhof, J., Pedersen, J.T., Fuglsang, A.T., Palmgren, M. Plasma Membrane H+-ATPase Regulation in the Center of Plant Physiology. Mol. Plant 2016, 9, 323–337.

37. Azad, A.K., Sawa, Y., Ishikawa, T., Shibata, H. Phosphorylation of Plasma Membrane Aquaporin Regulates Temperature-Dependent Opening of Tulip Petals. Plant Cell Physiol. 2004, 45, 608–617.

38. Yaneff, A., Vitali, V., Amodeo, G. PIP1 aquaporins: Intrinsic water channels or PIP2 aquaporin modulators? FEBS Lett. 2015, 589, 3508–3515.

39. Hachez, C., Zelazny, E., Chaumont, F. Modulating the expression of aquaporin genes in planta: A key to understand their physiological functions? Biochim. Biophys. Acta - Biomembr. 2006, 1758, 1142–1156.

40. Conde, A., Silva, P., Agasse, A., Conde, C., Gerós, H. Mannitol Transport and Mannitol Dehydrogenase Activities are Coordinated in Olea europaea Under Salt and Osmotic Stresses. Plant Cell Physiol. 2011, 52, 1766–1775.

41. Yang, L., Qin, L., Liu, G., Peremyslov, V. V; Dolja, V. V; Wei, Y. Myosins XI modulate host cellular responses and penetration resistance to fungal pathogens. Proc. Natl. Acad. Sci. 2014, 111, 13996–14001.

42. Sakurai, J., Ishikawa, F., Yamaguchi, T., Uemura, M., Maeshima, M. Identification of 33 Rice Aquaporin Genes and Analysis of Their Expression and Function. Plant Cell Physiol. 2005, 46, 1568–1577.

43. Verdoucq, L., Rodrigues, O., Martinière, A., Luu, D.T., Maurel, C. Plant aquaporins on the move: reversible phosphorylation, lateral motion and cycling. Curr. Opin. Plant Biol. 2014, 22, 101–107.

44. Qiu, J., McGaughey, S.A., Groszmann, M., Tyerman, S.D., Byrt, C.S. Phosphorylation influences water and ion channel function of AtPIP2;1. Plant. Cell Environ. 2020, 43, 2428– 2442.

45. Kawa, D., Testerink, C. Regulation of mRNA decay in plant responses to salt and osmotic stress. Cell. Mol. Life Sci. 2017, 74, 1165–1176.

46. Duque, P. A role for SR proteins in plant stress responses. Plant Signal. Behav. 2011, 6, 49–54.

47. Kong, X., Ma, L., Yang, L., Chen, Q., Xiang, N., Yang, Y., Hu, X. Quantitative Proteomics Analysis Reveals That the Nuclear Cap-Binding Complex Proteins Arabidopsis CBP20 and CBP80 Modulate the Salt Stress Response. J. Proteome Res. 2014, 13, 2495–2510.

48. Papp, I., Mur, L.A., Dalmadi, A., Dulai, S., Koncz, C. A mutation in the Cap Binding Protein 20 gene confers drought tolerance to Arabidopsis. Plant Mol. Biol. 2004, 55, 679–686.

49. Li, T., Gonzalez, N., Inzé, D., Dubois, M. Emerging Connections between Small RNAs and Phytohormones. Trends Plant Sci. 2020, 25, 912–929.

50. Rosa, M., Prado, C., Podazza, G., Interdonato, R., González, J.A., Hilal, M., Prado, F.E. Soluble sugars--metabolism, sensing and abiotic stress: a complex network in the life of plants. Plant Signal. Behav. 2009, 4, 388–393.

51. Yang, S.S., Zhai, Q.H. Cytosolic GAPDH: a key mediator in redox signal transduction in plants. Biol. Plant. 2017, 61, 417–426.

52. Kim, S.-C., Guo, L., Wang, X. Nuclear moonlighting of cytosolic glyceraldehyde-3-phosphate dehydrogenase regulates Arabidopsis response to heat stress. Nat. Commun. 2020, 11, 3439.

53. Kim, S.-C., Guo, L., Wang, X. Nuclear moonlighting of cytosolic glyceraldehyde-3-phosphate dehydrogenase regulates Arabidopsis response to heat stress. Nat. Commun. 2020, 11, 3439.

54. Wang, X., Cai, X., Xu, C., Wang, Q., Dai, S. Drought-Responsive Mechanisms in Plant Leaves Revealed by Proteomics. Int. J. Mol. Sci 2016, 17, 1.

55. Piattoni, C.V; Ferrero, D.M.L., Dellaferrera, I., Vegetti, A., Iglesias, A.Á. Cytosolic Glyceraldehyde-3-Phosphate Dehydrogenase Is Phosphorylated during Seed Development. Front. Plant Sci. 2017, 8, 522.

56. Chávez-Bárcenas, A.T., Valdez-Alarcón, J.J., Martínez-Trujillo, M., Chen, L., Xoconostle-Cázares, B., Lucas, W.J., Herrera-Estrella, L. Tissue-Specific and Developmental Pattern of Expression of the Rice sps1 Gene. Plant Physiol. 2000, 124, 641 LP – 654.

57. Toroser, D., Huber, S.C. Protein phosphorylation as a mechanism for osmotic-stress activation of sucrose-phosphate synthase in spinach leaves. Plant Physiol. 1997, 114, 947–955.

58. Zhen, S., Deng, X., Zhang, M., Zhu, G., Lv, D., Wang, Y., Zhu, D., Yan, Y. Comparative Phosphoproteomic Analysis under High-Nitrogen Fertilizer Reveals Central Phosphoproteins Promoting Wheat Grain Starch and Protein Synthesis. Front. Plant Sci. 2017, 8, 67.

59. Vener, A. V Environmentally modulated phosphorylation and dynamics of proteins in photosynthetic membranes. Biochim. Biophys. Acta - Bioenerg. 2007, 1767, 449–457.

60. Chen, Y.-E., Cui, J.-M., Su, Y.-Q., Zhang, C.-M., Ma, J., Zhang, Z.-W., Yuan, M., Liu, W.-J., Zhang, H.-Y., Yuan, S. Comparison of phosphorylation and assembly of photosystem complexes and redox homeostasis in two wheat cultivars with different drought resistance. Sci. Rep. 2017, 7, 12718.

61. Longoni, P., Douchi, D., Cariti, F., Fucile, G., Goldschmidt-Clermont, M. Phosphorylation of the Light-Harvesting Complex II Isoform Lhcb2 Is Central to State Transitions. Plant Physiol. 2015, 169, 2874–2883.

62. Riché, A., Lefebvre-Legendre, L., Goldschmidt-Clermont, M. A key role for phosphorylation of PsbH in the biogenesis and repair of photosystem II in Chlamydomonas. bioRxiv 2019, 754721.

63. Guidi, L., Lo Piccolo, E., Landi, M. Chlorophyll Fluorescence, Photoinhibition and Abiotic Stress: Does it Make Any Difference the Fact to Be a C3 or C4 Species? Front. Plant Sci. 2019, 10, 174.

64. Qiu, J., Hou, Y., Wang, Y., Li, Z., Zhao, J., Tong, X., Lin, H., Wei, X., Ao, H., Zhang, J. A Comprehensive Proteomic Survey of ABA-Induced Protein Phosphorylation in Rice (Oryza sativa L.). Int. J. Mol. Sci. 2017, 18, 60.

65. Schwartz, D., Gygi, S.P. An iterative statistical approach to the identification of protein phosphorylation motifs from large-scale data sets. Nat. Biotechnol. 2005, 23, 1391–1398.

66. Bigeard, J., Hirt, H. Nuclear Signaling of Plant MAPKs. Front. Plant Sci. 2018, 9, 469.

67. Sinha, A.K., Jaggi, M., Raghuram, B., Tuteja, N. Mitogen-activated protein kinase signaling in plants under abiotic stress. Plant Signal. Behav. 2011, 6, 196–203.

68. van Wijk, K.J., Friso, G., Walther, D., Schulze, W.X. Meta-Analysis of Arabidopsis thaliana Phospho-Proteomics Data Reveals Compartmentalization of Phosphorylation Motifs. Plant Cell 2014, 26, 2367 LP – 2389.

69. Shinozawa, A., Otake, R., Takezawa, D., Umezawa, T., Komatsu, K., Tanaka, K., Amagai, A., Ishikawa, S., Hara, Y., Kamisugi, Y., et al. SnRK2 protein kinases represent an ancient system in plants for adaptation to a terrestrial environment. Commun. Biol. 2019, 2, 30.

70. Vilela, B., Pagès, M., Riera, M. Emerging roles of protein kinase CK2 in abscisic acid signaling. Front. Plant Sci. 2015, 6, 966.

